# Protein Embeddings Predict Binding Residues in Disordered Regions

**DOI:** 10.1101/2024.03.05.583540

**Authors:** Laura R. Jahn, Céline Marquet, Michael Heinzinger, Burkhard Rost

## Abstract

The identification of protein binding residues helps to understand their biological processes as protein function is often defined through ligand binding, such as to other proteins, small molecules, ions, or nucleotides. Methods predicting binding residues often err for intrinsically disordered proteins or regions (IDPs/IDPRs), often also referred to as molecular recognition features (MoRFs). Here, we presented a novel machine learning (ML) model trained to specifically predict binding regions in IDPRs. The proposed model, IDBindT5, leveraged embeddings from the protein language model (pLM) ProtT5 to reach a balanced accuracy of 57.2±3.6% (95% confidence interval). Assessed on the same data set, this did not differ at the 95% CI from the state-of-the-art (SOTA) methods ANCHOR2 and DeepDISOBind that rely on expert-crafted features and evolutionary information from multiple sequence alignments (MSAs). Assessed on differ data, methods such as SPOT-MoRF reached higher MCCs. IDBindT5’s SOTA predictions are much faster than other methods, easily enabling full-proteome analyses. Our findings emphasize the potential of pLMs as a promising approach for exploring and predicting features of disordered proteins. The model and a comprehensive manual are publicly available at https://github.com/jahnl/binding_in_disorder.

## Introduction

Intrinsically disordered proteins (<u>IDPs</u>) [1], or local regions in these (<u>IDPRs</u>) are characterized by regions of amino acids absent of regular secondary [2] or tertiary structure [3-6], exhibiting unique properties that enable them to maintain domain separation, facilitate interactions, and offer increased propensity for post-translational modifications, such as phosphorylation [3]. IDPs are involved in signal transduction, regulation, and specific binding with various partners depending on the cellular environment, including metal ions, inorganic molecules, small organic molecules, and other biomolecules [3, 7]. Understanding binding in IDPs may provide important insights, including potential drug targeting [3] and may offer opportunities to engineer proteins and to design biomaterials for diverse scientific and medical purposes [8].

The experimental identification of binding residues in IDPs remains challenging [9]. This yielded an avalanche of *in silico* methods for IDPs as tools geared toward well-structured proteins often fail on disordered proteins [9]. IDPs and well-structured proteins are not antipodes, instead there is a spectrum of transitions, including partial disorder, hybrid proteins, or conditional disorder, wherein environmental conditions may change structure [3]. The latter can occur when IDPRs change to ordered regions upon binding in segments known as molecular recognition features (<u>MoRFs</u>). Identifying semi-disordered regions is useful to locate functional IDPRs, which in turn enables Machine Learning (<u>ML</u>) tools to successfully identify MoRFs [10, 11]. However, the inherent complexity of disorder transition complicates the curation of gold standards for ML [3, 12].

The Critical Assessment of protein Intrinsic Disorder (<u>CAID</u>) project has been providing sustained evaluations [13]. In 2018, *ANCHOR2* performed best [13, 14] by using biophysics-based energy functions balancing disorder tendency and interaction [14]. Two years later, methods outperformed ANCHOR2 [15, 16] through using homology-based inference (<u>HBI</u>) [12, 13], evolutionary information from multiple sequence alignments (<u>MSAs</u>) and expert-crafted, knowledge-based features as input for ML [12, 14, 15]. However, for disordered proteins, this information is often unavailable [17], and it has been shown that it may be suboptimal to apply MSAs and hand-crafted features to IDPs and IDPRs [9].

Protein language models (<u>pLMs</u>) present a novel approach for representing protein sequences without explicitly using MSAs. Drawing inspiration from Natural Language Processing (<u>NLP</u>), pLMs utilize the sequential order of input to learn the underlying principles of sequence features in an unsupervised and hypothesis-free manner [18, 19]. The advantage of this approach lies in its independence from manually validated data, allowing for substantial scalability of generic Large Language Models (<u>LLMs</u>) [18]. Prominent examples for pLMs are ProtT5 [18], ESM-2 [20], SeqVec [21] and Progen [22], all successfully applying transformer architectures on protein sequences. Since the introduction of transformer-based pLMs, it has been shown that the protein embeddings, i.e., numeric representations generated with such pre-trained models (practically the values from the last hidden layers [18]), are predictive of many aspects of protein structure and function, performing *on par* or outperforming MSA-based methods while only using sequence as input [18, 20, 23-31].

Inspired by the success of MSA-free pLM embeddings particularly for binding residue prediction [23], we introduce *<u>IDBindT5</u>* (Fig. 1), a novel method predicting binding residues in disordered regions (IDPRs) from single sequence input. We trained IDBindT5 on carefully curated and redundancy-reduced data sets derived from *MobiDB* [32], which collects data on protein disorder and interactions from several other databases, such as *DisProt* [7] and *IDEAL* [33]. Leveraging ProtT5 embeddings as input, IDBindT5 predicts binary scores indicating whether a residue in an IDPRs is likely to bind.

**Fig. 1:**
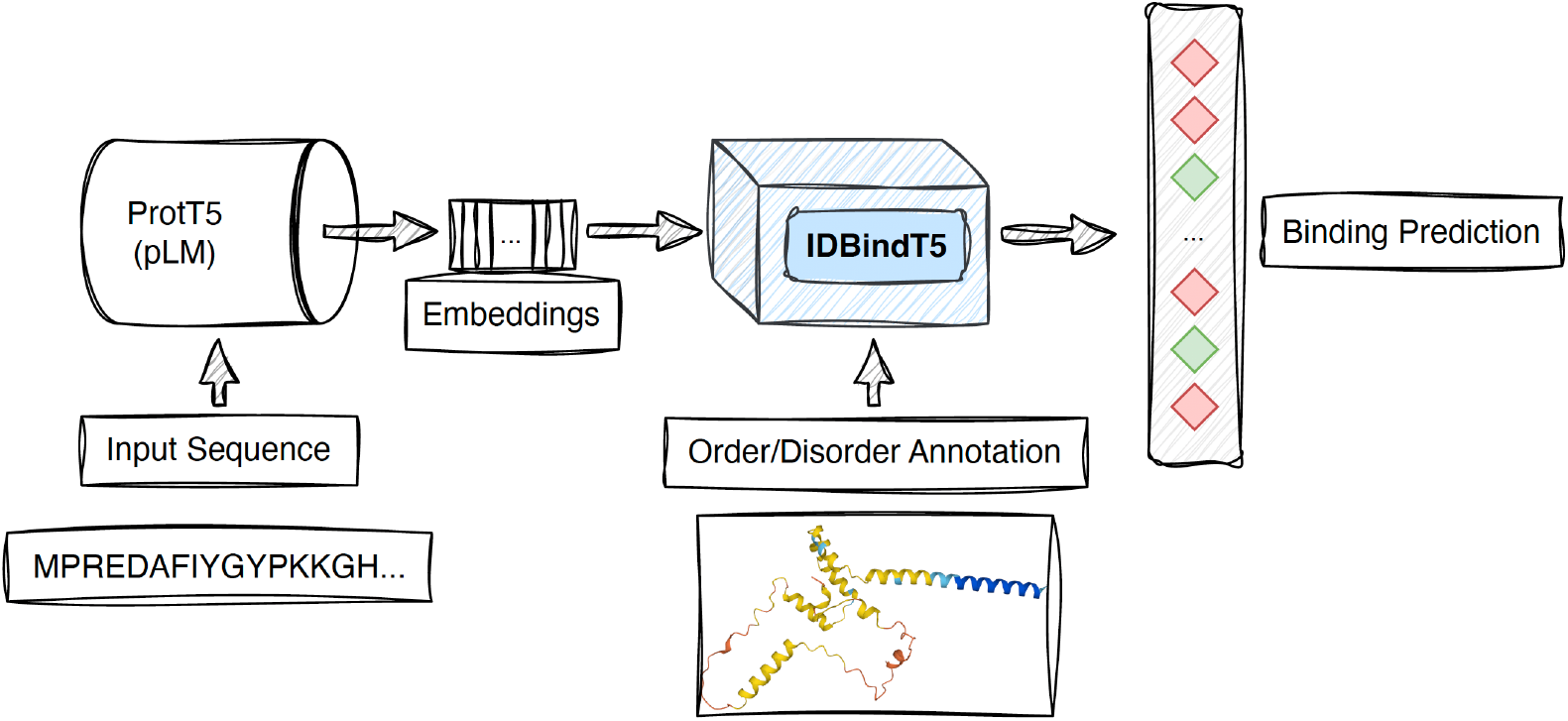
Workflow of IDBindT5. To predict whether or not a residue in a disordered region (IDPR) is binding, *IDBindT5* takes numerical representations of a protein sequence (embeddings) generated by the protein language model (pLM) ProtT5 [18] accompanied by either predicted or experimental per-residue annotation of (dis-)order as input. *IDBindT5* avoids dramatic explosion of free parameters and overfitting through a relatively simple feedforward network (FNN, single hidden layer). The input consists of a binary vector for disorder/order (either from prediction or annotation; dimension L × 1, with L being the number of residues in a given protein), and the generated embeddings are of shape L × m (m depends on specific pLM, for ProtT5 m = 1024). The output produced by *IDBindT5* is of shape L × 1, representing a per-residue binding prediction.

## Results and Discussion

We propose the novel machine learning (ML) model IDBindT5, which predicts binding residues in intrinsically disordered protein regions (IDPRs). The feed-forward neural network (FNN) with a single hidden layer takes embeddings from the protein language model (pLM) ProtT5 [18] and disorder annotation as input and outputs a binary per-residue binding prediction.

### Choice of pLM relevant but not critical

We compared several pLMs as input embeddings for the downstream prediction task of binding within IDPRs. Comparing “identical” ML architectures (only differing in the dimensionality of the input pLMs), the model based on ProtT5 [18] slightly outperformed the one based on ESM-2 [20]: MCC(IDBindT5) = 0.210 ± 0.019 (95% CI, confidence interval) vs. MCC(FNN_all_ESM2) = 0.179 ± 0.019 (Supplementary Fig. SOM_F2 and Table SOM_T3 in the *Supporting Online Material (SOM)*). This difference was numerically better, but not statistically significant within the 95% CI (confidence interval, i.e., at ±1.96 standard errors). Other performance measures largely confirmed the same trend (Supplementary Table SOM_T3). Thus, the choice of the pLM appeared, overall, not critical, confirming similar results for conservation prediction [26]. This also underscored the robustness of embeddings from different pLMs for subsequent supervised training.

### Embeddings allow for simple downstream architecture

We developed several ML models. On the validation set, two tied for best performance: a CNN (Convolutional Neural Network) trained on disordered regions only (dubbed CNN_disorder) and IDBindT5 (Supplementary Fig. SOM_F2 and Table SOM_T3), an FNN (Feedforward Neural Network with single hidden layer) trained on both ordered and disordered residues. As the CNN can capture more information about the local residue environment than an FNN, this result suggested that the ProtT5 embeddings already captured that information. To reduce the odds of over-fitting and limit the runtime/energy consumption, we favored the FNN as the final model.

In contrast to a CNN, which was balanced on a per-protein level, an FNN also provided the option to incorporate per-residue level balance through over-or under-sampling on the per-residue level. Consequently, the CNNs differed more on the validation set between the two training datasets than the FNNs. Training a CNN on all residues as opposed to only on disordered residues reduced the F1 (Eqn. 6) by about 10%, for the FNNs, the corresponding difference remained below 1% (Supplementary Table SOM_T3). Since CNN and FNN performed similarly on the validation set, we tested their consensus combination (average over raw output and optimization of threshold). Performance increased numerically, but not at levels of statistical significance (Supplementary Table SOM_T4).

### IDBindT5 reached SOTA

After the above steps decided on the best model (IDBindT5) according to the validation set, we estimated performance through the test set (Fig. 2, Supplementary Table SOM_T4). Compared to the validation set, test set estimates of precision and recall were both significantly lower (Supplementary Tables SOM_T3 and SOM_T4), while balanced accuracy (Eqn. 5) and MCC (Eqn. 7) remained within the 95% CI of each other (respective p-values via Welch’s t-test: 0.580 and 0.790). The decrease in estimates for precision and recall from validation to test set were similar for the random baseline. Neither was “compensated” for by a change in the corresponding values for negatives. IDBindT5 clearly outperformed the random baseline.

**Fig. 2:**
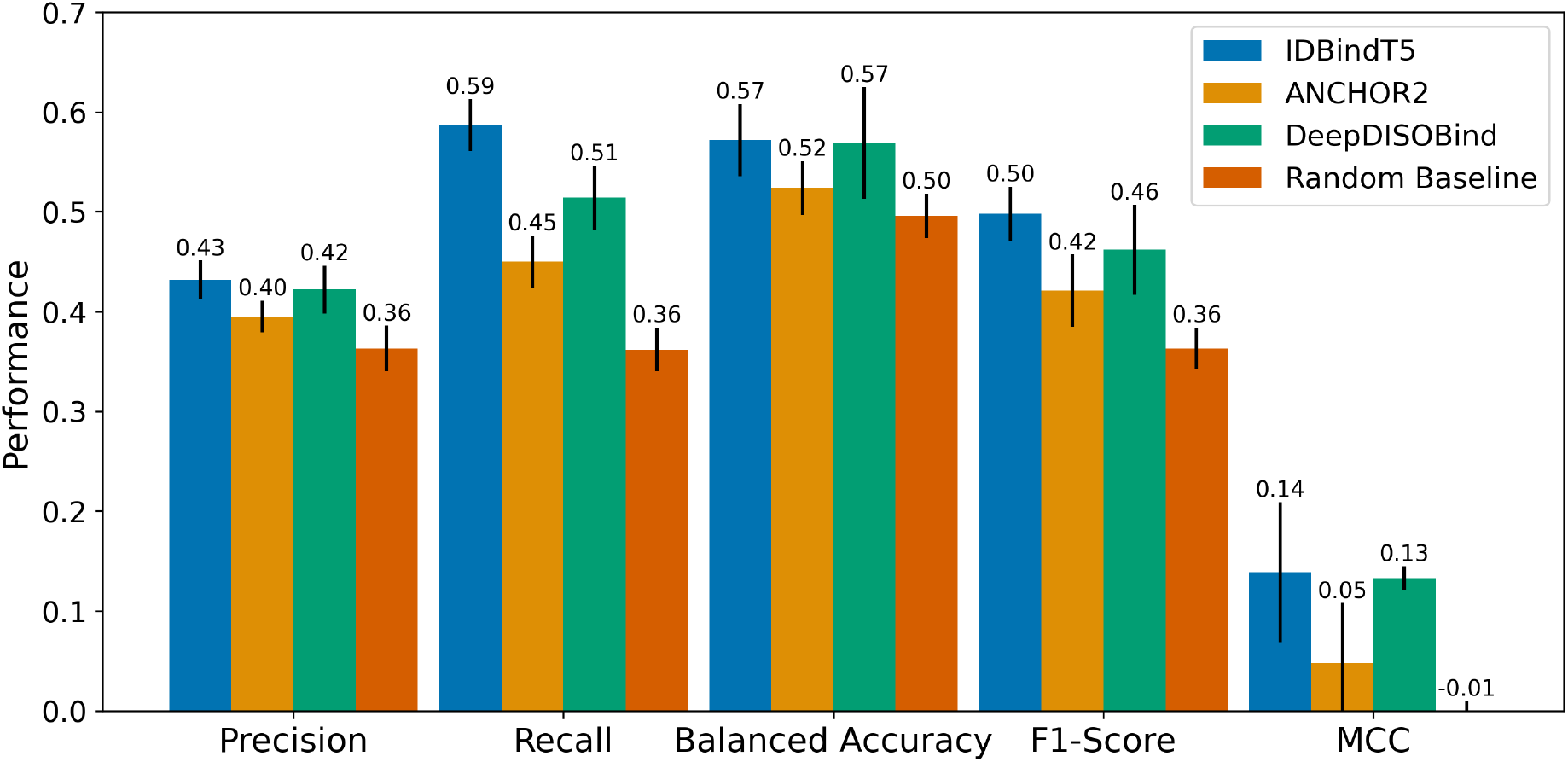
Test set performance. Methods to predict binding regions in IDPRs (intrinsically disordered regions): *IDBindT5* (here), SOTA methods *ANCHOR2* [14] and *DeepDISOBind* [34], and a *Random baseline* (frequency of 38.8% positives and 61.2% negatives). Data set: test set (*Mobi195* with 195 proteins). The performance is captured by per-residue measures (scaled to [0,1]): precision (Eqn. 1), recall (Eqn. 2), balanced accuracy (Eqn. 5), F1-Score (Eqn. 6) and MCC (Eqn. 7). Error bars reflect the 95% confidence interval (CI) of 1.96*standard errors. Only IDBindT5 is significantly better than the random baseline for all measurements. While it also reaches the highest numerical values for all measures, not all differences are significant at 95% CI. Noteworthy is the superior recall value at top precision (more measures in Supplementary Table SOM_T4).

In comparison to the SOTA models *ANCHOR2* [14] and *DeepDISOBind* [34], IDBindT5 performed favorably in our hands (Fig. 2, Supplementary Table SOM_T4). However, while IDBindT5 was numerically best in terms of balanced accuracy and MCC, no method stood out in a Welch test [35] (Supplementary Table SOM_T7). Both recall and precision of IDBindT5 exceeded those for the two SOTA methods (Supplementary Table SOM_T4), again without consistent statistical significance (Supplementary Table SOM_T7).

The number of consecutive residues predicted as binding by IDBindT5 (mean: 4, median: 1) and DeepDISOBind (mean: 18, median: 1) mostly comprised single residues, while experimentally determined binding regions in the test set constituted much longer stretches (mean: 74, median: 41; Supplementary Figs. SOM_F5-7). ANCHOR2’s predictions were closer to the experimental profiles than the other two methods (mean: 39, median: 20).

The test set presented in this work was redundancy reduced against the training sets of all evaluated methods based on sequence and structure. While we could not consider the time-based split of the first Critical Assessment of protein Intrinsic Disorder (CAID) benchmark [13], the deadline for CAID2 [15] (November 2022) was applicable to our training set (cutoff October 2022). On the CAID2 benchmark, IDBindT5 performed similarly to the level in our test set compared to the other SOTA methods with IDBindT5 and DeepDISOBind reaching numerically higher performance than ANCHOR2, but not differing at the 95% CI. (Fig. 3).

**Fig. 3:**
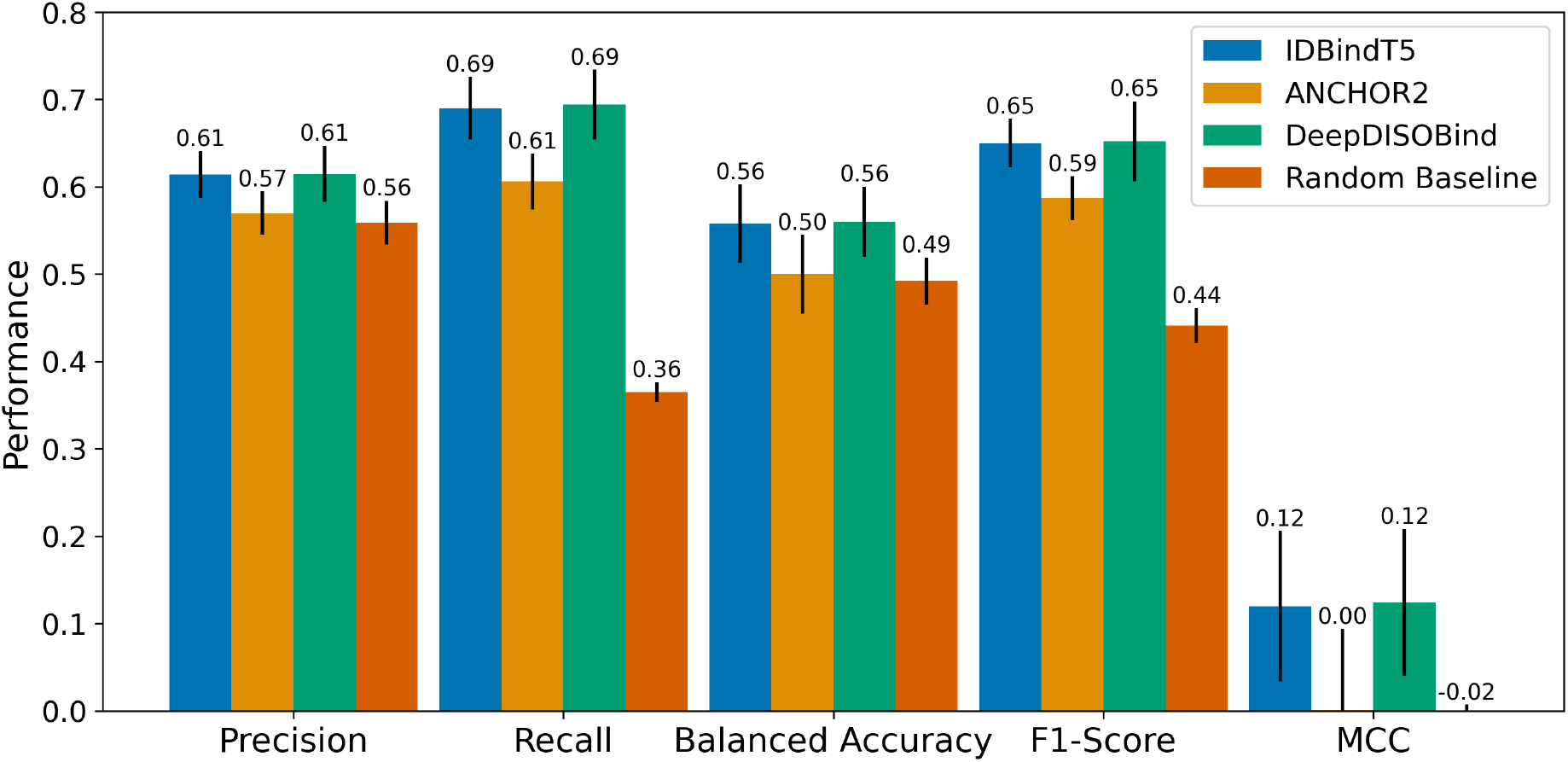
CAID2 binding benchmark. The Critical Assessment of protein Intrinsic Disorder round 2 (CAID2 [15]) assessed methods to predict binding regions in IDPRs (intrinsically disordered protein regions). Here, we showed *IDBindT5* (method introduced in this paper) along with the SOTA methods *ANCHOR2* [14] and *DeepDISOBind* [34], and a *Random baseline* (randomly assigning frequency of positives and negatives). Data set: CAID2 binding set with 78 proteins. The performance is captured by the four per-residue measures (scaled to [0,1]): precision (Eqn. 1), recall (Eqn. 2), balanced accuracy (Eqn. 5), F1-Score (Eqn. 6), and MCC (Eqn. 7). Error bars reflect the 95% confidence interval (CI) of 1.96*standard errors. While IDBindT5 and DeepDISOBind reach higher numerical performance than ANCHOR2, not all differences are significant at 95% CI.

Given the small sizes of both our test set (195 proteins) and the CAID2 binding benchmark (78 proteins), none of the SOTA methods consistently improved over the random baseline (randomly assigning positives and negatives based on frequency of experimental annotations) at levels that were statistically significant (Fig. 2, Fig. 3). Due to the difference in training data and redundancy-reduction used for the development of other SOTA methods, we restricted comparison to the two SOTA methods ANCHOR2 and DeepDISOBind evaluated on CAID1 [13]. Future analyses including other SOTA tools from CAID2 [15] and tools such as SPOT-MoRF [10] which were not evaluated on CAID but have been shown to outperform ANCHOR2 [14], may yield additional insights.

### IDBindT5 performance with predicted disorder data remains high

An ablation study analyzing the input feature importance of IDBindT5 (1024-dimensional ProtT5 embedding vector and one-dimensional binary disorder annotation; Supplementary Fig. SOM_F8) suggested that the input unit disorder carried more weight than any other single dimension. Surprisingly, it contributed only ∼50% more than the next strongest single unit, and at least 19 other units contributed half as much to the final decision (Supplementary Fig. SOM_F8).

We also evaluated the performance on the test set with disorder labels predicted by the SOTA methods SETH (pLM-based) [36] and AlphaFold2-disorder (structure-based) [17] (Supplementary Table SOM_T5). The disorder predictions on our test set remained somehow limited (MCC = 0.324, and 0.432 for SETH and AlphaFold2-disorder, respectively; Supplementary Table SOM_T5). The binding predictions of IDBindT5 using those faulty predictions as input rather than the experimental annotations remained surprisingly stable considering levels of precision below 40% (Supplementary Table SOM_T5). Only recall, and consequently the F1 score, dropped levels reaching statistical significance to 39.5±3.0% (SETH input) or 42.3±2.8% (AlphaFold2 input). All other performance metrics remained within the CI of the performance with curated disorder data (Supplementary Table SOM_T4).

### IDBindT5 good and fastest

Users want methods to be as good as possible, and developers making their methods available on webservers want them to be energy-aware, i.e., to consume as few resources as possible. Thus, we compared the three best methods on *Mobi11k*, a subset of 11,114 human proteins (median length 489 residues; see *Methods*) using the same machine (CPU: 2x Intel Xeon Gold 6248, 400 GB RAM, 20 cores; GPU: 4x Quadro RTX 8000, 46 GB vRAM). The slowest method, DeepDISOBind needed 33 hours and 22 minutes, the fastest, ANCHOR2 took 21 minutes. Our method, IDBindT5, completed predictions within 2 hours and 10 minutes, providing a balance between predictive performance and runtime. To analyze single sequence runtime, we used a consumer grade laptop (CPU: Intel Core i7, 16 GB RAM, 4 cores; GPU: NVIDIA GeForce MX350, 9.9 GB vRAM). Averaged over 10 runs for a median length disordered protein in our training data (399 residues), IDBindT5 took around 2 seconds on CPU. There was no advantage when using GPU for single sequence prediction, but it reduced the runtime on the dataset Mobi195 by 35% (see Supplementary Table SOM_T8 for all runtime measurements and conditions).

### Wanted: additional experimental data on binding in disordered regions

Although bigger AI models are not always better in computational biology [18], without models with substantially more free parameters than three decades ago, neither pLMs nor breakthroughs on the level of AlphaFold2 [37] would have been feasible. Although the past rule-of-thumb of 10 of the minority-class samples (here the positives accounting for 38% of all points) per free parameter are more often ignored by orders of magnitude than not in the era of deep learning, modern AI is simply data hungry. In fact, this is the reason why generalized pLMs are so successful: they accumulate over 50-times the amount of data as the entire Wikipedia of the English language for Natural Language Processing [18, 21]. IDPs and IDPRs clearly do not fall into this category. In fact, pLM embeddings help through their power in transfer learning, i.e., by partially *understanding the* language *of life as written in protein sequences* [21], they allow the development of relatively simple ML models to predict disorder without using too much data [31]. Our application of pLMs to predict binding in IDPRs clearly pushed the envelope.

The inherent complexities of molecular interactions in IDPRs render experimental annotations of protein-protein interactions in such regions exceptionally daunting. Therefore, available data to train ML methods is sparse and necessitated strategic compromises for the development of IDBindT5. While IDBindT5 provides accurate and fast SOTA per-residue binding predictions in disordered regions, it does not distinguish between the type of binding and while it maintains SOTA performance with predicted disorder as input, it depends on the overall quality of such predictions (Supplementary Table SOM_T5). Although other methods are able to predict DNA and RNA binding and protein hubs, performance is often very limited with a large performance gap compared to binding site prediction in well-structured regions [14, 15, 34] which in itself suffers greatly from a lack of reliable experimental data [23, 38]. We decided to maximize the available data by using binary annotations (binding/non-binding).

The database MobiDB [32] collects curated disorder and interaction information from several other databases, including IDEAL [33], UniProt [39], DIBS [40], ELM [41], MFIB [42], and DisProt [7]. As MobiDB provides 78% more IDPs with curated information than DisProt (2853 vs. 1605, as of September 2022), it was the natural choice to train an ML method with the focus of generalizability. However, the type of the binding partner when downloading entries from MobiDB via its application programming interface (API) is not listed, so training a multilabel model predicting ligand types was not possible.

Another compromise imposed by the data shortage was that the models were evaluated only on disordered regions. This choice was driven by the large data imbalance between negatives (anything in order plus non-binding residues in disorder, 93%) and positives (binding residues in disorder, 7%) (Supplementary Table SOM_T1). This imbalance naturally favors methods that mainly distinguish ordered from disordered residues, even when considering metrics that take class imbalance into account. In order to avoid a bias towards these models, we only evaluated the ability to predict binding in disordered residues. Consequently, our method numerically improved over the SOTA, but at the cost of requiring additional input data, disorder annotations, which the reference methods either do not need or predict by themselves during runtime.

The challenges encountered in developing a method to predict binding residues in IDPRs underscored the critical need for improved annotations in protein-protein interactions, particularly within disordered regions. Unfortunately, we cannot foresee whether a data duplication will significantly improve performance. But overcoming some of today’s data limitations might set the stage for future advancements that promise to deepen our understanding of molecular interactions within disordered regions, opening new avenues for research and discovery in the realm of structural and functional biology.

## Conclusions

We presented a new method, dubbed IDBindT5, for predicting whether a residue in an intrinsically disordered protein region (IDPR) binds to any ligand. The method achieved a balanced accuracy of 57.2±3.6%, an MCC of 0.139±0.07, and a precision of 43.2±1.9% at a recall of 58.7±2.6%. IDBindT5 reached similar levels as the state-of-the-art methods ANCHOR2 [14] and DeepDISOBind [34] (Fig. 2) albeit requiring neither alignments nor a complex ML architecture. Key to this is that IDBindT5 operates solely on embeddings from single sequences generated by the pre-trained pLM ProtT5 [18] and a disorder annotation as input. Replacing annotated by predicted disorder (SETH [36] and AlphaFold2-disorder [17]) reduced performance substantially (Supplementary Table SOM_T4) although this reduction was not even statistically significant within a confidence interval of 95% (± one standard error). In terms of prediction speed our model was over 15 times faster than DeepDISOBind, enabling many predictions even on common notebook hardware and requiring less energy. While overall, binding in IDPRs continues to be challenging, the use of pLM embeddings as input appears to be an approach worth building upon. IDBindT5, along with a comprehensive manual, is publicly available at https://github.com/jahnl/binding_in_disorder.

## Methods

### Datasets

We used four datasets: one for training and validation of our new method IDBindT5, two for testing and performance evaluation and comparison to the state-of-the-art (SOTA) methods ANCHOR2 [14] and DeepDISObind [34], and one for prediction speed assessment.

#### Train: Mobi2k and Test: Mobi195

For ease of comparisons, we relied upon MobiDB [32] (Oct. 17, 2022). To ensure the reliability of annotated proteins for the task of predicting the potential interaction between disordered residues and binding partners, we only selected proteins with manually curated disorder and binding annotations. The resulting data set comprised 2,816 proteins with 188,940 disordered residues. Of these disordered residues, 117,122 (62%) were categorized as negatives (disordered, non-binding), and 71,818 (38%) as positives (disordered, binding). We excluded the protein titin (Q8WZ42 with 34,350 residues) to manage memory consumption (only proteins with over 10k (10,000) residues).

The following meticulous redundancy reduction steps ensured an unbiased and robust assessment of performance. First, we randomly selected 200 positive (at least one binding region in disorder) and 200 negative proteins (no binding region in disorder) from the MobiDB dataset as a test set. To minimize sequence and structural similarity within the test set, we applied MMSeqs2 and UniqueProt [43-45]. First, we ran MMseqs2 [44] with default settings (e-value: 10, sensitivity: 7.5, minimum alignment length: 11) and calculated the HVAL (HSSP-value [43]). We reduced redundancy at HVAL=0 (corresponding to, e.g., <20% pairwise sequence identity (PIDE) for pairs of proteins aligned over 250 residues) for both positives and negatives, leaving 168 positives and 151 negatives. Next, we reduced the remaining negatives against the positives using the same procedure, resulting in 132 negatives. To facilitate comparisons with ANCHOR2 [14] and DeepDISObind [34], we redundancy-reduced the test set with respect to their training sets, resulting in 98 positives and 115 negatives.

The proteins removed this way due to overlap with the development sets of other methods, we added to our training set, giving 2,502 training set proteins. Next, we redundancy-reduced positives and negatives of the training set through CD-HIT [46] at 40% PIDE resulting in 777 positives and 1282 negatives. As before, we next redundancy-reduced the negatives against the positives (CD-HIT <40%), leaving 1,176 negatives. For added security against data leakage, we again redundancy reduced the resulting training set against the test set using MMSeqs2-UniqueProt [43-45] at HVAL<0, resulting in 1762 proteins (679 positives and 1083 negatives) in the training set.

Lastly, to balance the positives-to-negatives ratio in training and testing set, we excluded 18 positives (with 16,567 residues) from the test set and moved those to the training set, leading to the final training set, dubbed **<u>Mobi2k</u>**, with 1,780 proteins (697 positives, 1083 negatives, in disorder: 64,228 binding residues, 101,381 non-binding residues, Supplementary Table SOM_T1). Next, we randomly split the training set into five equally sized folds for five-fold cross-validation (CV). We used two versions of this set for training: one with all residues, and one consisting of only residues in annotated disordered regions. The final test set, dubbed **<u>Mobi195</u>**, consisted of 195 proteins (80 positives, 115 negatives, in disorder: 6,657 binding residues, 11,644 non-binding residues, Supplementary Table SOM_T1). In each split, one fold served as validation set to optimize hyper-parameters, including the choice of the best method. We estimated the test performance only on the best model identified by the validation set, i.e., avoided the frequent mistake of reporting test results for more than one of our models, typically the one performing best on the test set. The number of (non-)binding proteins, and the number of (non-)binding residues of each validation set are listed in Supplementary Table SOM_T1.

#### CAID2 – binding benchmark set

The second round of the Critical Assessment of Intrinsic protein Disorder (CAID) challenge [15] is a community-based blind test to determine the SOTA disorder and binding prediction methods. The benchmarks are based on a time split (Nov. 20, 2022), extracting the difference between the public and private versions of the DisProt database [7]. The benchmark set for binding in IDPRs comprised of 78 proteins with 8,209 binding residues (12.2%), accessible via https://caid.idpcentral.org/challenge/results.

#### Mobi11k – speed assessment

To assess the runtime needed to apply our method, we generated a third, larger dataset. We filtered the human proteome in the MobiDB at <100% PIDE and we included sequences shorter than 5,000 residues with an annotation matching the sequence length and having at least 5% disordered residues predicted by MobiDB-lite [47]. This resulted in 11,114 human proteins with median sequence length of 489 residues.

### Input pLM embeddings

As input, we compared embeddings from the following non-fine-tuned pLMs: (1) ESM-2 with 3 billion (B) parameters [20] (based on BERT [48], trained on UniRef50 [39]), and (2) ProtT5-XL-U50 [18] with 3B parameters (for simplicity referred to as ProtT5; based on T5 [49], trained on BFD [50] and Uniref50 [39]). We extracted the per-residue embeddings from the last hidden layer of the models of dimension d=1280 for ESM-2 and d=1024 for ProtT5 × *L*, where L is the length of the protein sequence and d is the dimension of the hidden states/embedding space of ESM-2 or ProtT5. The embeddings were expanded by one dimension containing a binary indicator whether the residue is located in an IDPR based on the manually curated information from MobiDB [32].

### Prediction methods

As shown before [23-27], the rich information contained in embeddings allows to keep models for downstream prediction tasks relatively shallow (million(s) instead of billions of free parameters). We systematically combined the following architectures and parameters, creating different models: Architecture (Feed-forward Neural Networks (FNNs) and Convolutional Neural Networks (CNNs), balancing strategies (over-and under-sampling), training residue subsets (all residues of a protein or disordered only), number of layers, kernel size (for CNNs only), dropout probability, batch size and learning rate. All models were implemented in PyTorch [51], applied early stopping with a patience (i.e., number of epochs without improvement in loss on the validation set) of five and used the Adam [52] optimizer. Non-linearity was introduced by using the ReLU activation function after each hidden layer (except for the last hidden layer). As loss function, we applied the binary cross entropy loss. The training progress is visualized in Supplementary Fig. SOM_F3. The cutoff used to classify a residue as binding (output probability of the model > cutoff) or non-binding (output probability ≤ cutoff) was chosen individually for each model and fold, based on receiver operating characteristic (ROC) curves (Supplementary Fig. SOM_F1; precision-recall curves see Supplementary Fig. SOM_F4). The aim was to achieve a good precision while avoiding a large decrease in recall. All models exclusively input pLM embeddings and per-residue disorder annotations.

For each combination of parameters and architectures, we trained five models, one for each CV-fold. These were combined to an ensemble method by averaging their final output. Below, we describe a subset of the methods, chosen based on validation-set performance (for all methods see Supplementary Table SOM_T2).

#### IDBindT5

Two approaches performed best on the validation set: a CNN trained on disordered regions only (dubbed CNN_disorder) and IDBindT5 (Fig. 1), an FNN trained on both ordered and disordered residues (Supplementary Fig. SOM_F2, Table SOM_T3). Reducing the odds of over-fitting and limiting the runtime/energy consumption, we chose the FNN for IDBindT5 (input: ProtT5 (Fig. 1; 1025 units), hidden layer (612 units), and output layer (one unit), leading to a total of 1.7M (million) free parameters, trained with a batch size (here: number of residues) of 512, and a learning rate of 0.01). We trained the model using both ordered and disordered (“all”) residues (Supplementary Tables SOM_T1, SOM_T2). The FNN architecture was better suited for balancing the dataset on residue-level and given the unbalanced dataset, we decided to under-sample the non-binding residues within and outside of IDPRs. Due to the different distributions in the CV-folds, we applied a different ratio for each so that both subsets reach the same abundance as the binding residues for each fold. Based on the predictive performance on the validation set, the prediction cutoffs for the five models of the different CV-folds were chosen based on the respective ROC curves as 0.55, 0.40, 0.50, 0.35 and 0.30 (Supplementary Fig. SOM_F1 c). The model evaluated on the last CV fold had the best performance on the validation set and was thus selected to represent our final method – IDBindT5 (inference cutoff 0.30 with binding (output probability > 0.30) and non-binding (output probability ≤ 0.30)).

#### FNN_all_ESM2

We trained a model with the same architecture and training settings as IDBindT5 but switching the ProtT5 [18] embeddings for those of ESM-2 [20] with 3B parameters. Due to the larger vector size (2,560 input units for ESM-2), the free parameter count increased from 1.7M (ProtT5) to 9.8M (ESM-2). The cutoffs for the five models of the different folds were chosen as 0.4, 0.35, 0.45, 0.4 and 0.35, respectively (Supplementary Fig. SOM_F1 d).

#### Additional Baselines

As **<u>random baseline</u>** we only considered the frequency of positives in the training set. Thus, we randomly predicted 38.8% of the disordered residues as binding. In addition to a random baseline, we introduced a baseline trained on classical biophysical amino acid (AA) features, dubbed **<u>AAindex_disorder</u>**. *AAindex1* is a database of numerical properties of single amino acids, derived from literature [53]. Instead of the features of the protein embedding, a vector of all 566 associated AAindex1 values was assigned to each residue. The model was then trained based on the configuration of the model *CNN_disorder* (Supplementary Table SOM_T2, Supplementary Fig. SOM_F1 f).

### Performance measures

The following standard measures assessed performance (<u>TP</u> (true positives): correctly predicted as binding; <u>FP</u> (false positives): incorrectly predicted as binding; <u>TN</u> (true negatives): correctly predicted as non-binding; <u>FN</u> (false negatives): incorrectly predicted as non-binding). The resulting 2x2 confusion matrix enabled the computation of all non-statistical scores used for evaluation, namely:

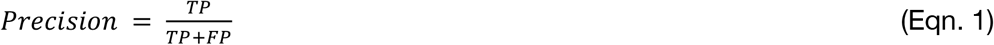

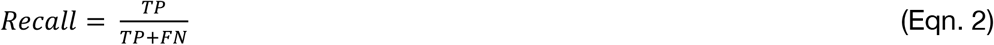

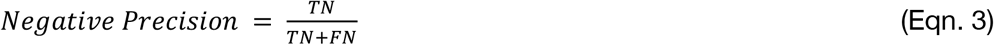

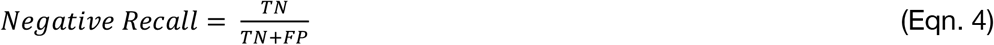

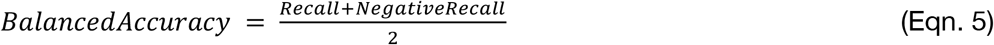

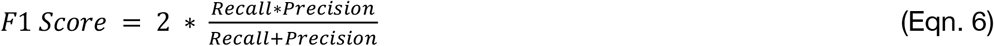

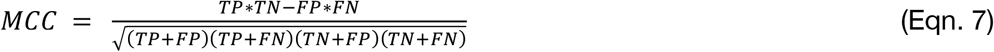

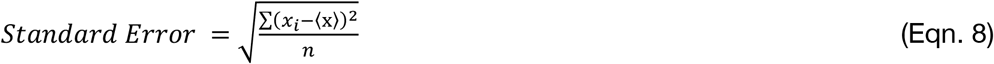

Standard boot-strapping [54] turned these metrics (Eqn. 1-7) into statistical estimates, applying each to batches of 100 disordered residues. To avoid protein length affecting the proportional impact of a prediction, we averaged over the resulting distribution. We estimated the standard error by bootstrapping 1k batches (Eqn. 8), then we applied a 95% confidence interval (CI) [54, 55]. The difference between distributions, whose 95% CIs did not overlap, was considered as statistically significant. To compare models whose 95% CIs overlapped and consequently showed no significant difference, we performed Welch’s t-test which does not assume equal population variance [35, 55].

#### Comparison to SOTA

As state-of-the-art methods (SOTA), we compared IDBindT5 to ANCHOR2 [14] and DeepDISOBind [34]; both stood out at CAID [13, 56]. We tapped into the webserver of each resource to obtain predictions for all proteins in our test set. While ANCHOR2 predicts only protein-protein interactions, DeepDISOBind also predicts interactions with nucleic acids [34].

## Abbreviations used

AA: Amino Acid
AI: Artificial Intelligence
API: Application Programming Interface,
BFD: Big Fantastic Database
CAID: Critical Assessment of protein Intrinsic Disorder
CNN: Convolutional Neural Network
CV: Cross Validation
FN: False Negative
FNN: Feed-forward Neural Network
FP: False Positive
HBI: Homology-Based Inference
HVAL: HSSP Value
IDP: Intrinsically Disordered Protein
IDPR: Intrinsically Disordered Protein Region
LLM: Large Language Models
MCC: Matthews Correlation Coefficient
ML: Machine Learning
MoRF: molecular recognition features,
MSA: Multiple Sequence Alignment
NLP: Natural Language Processing
NN: Neural Network
PIDE: Pairwise sequence Identity
pLM: protein Language Model
ROC: Receiver Operating Characteristic
SOM: Supporting Online Material
TN: True Negative
TP: True Positive

## Data availability

The source code, data sets, the trained models and a detailed manual on how to use our prediction tool are publicly available in the project’s GitHub repository (https://github.com/jahnl/binding_in_disorder). ProtT5 embeddings can be generated using the *bio_embeddings* [18] pipeline or the Jupyter Notebook provided in the ProtTrans GitHub repository (https://github.com/agemagician/ProtTrans). Disorder annotations can be replaced by predictions from the LambdaPP webserver [57].

## Acknowledgements

Thanks to Nikita Kugut, Inga Weise (both TUM) and Tim Karl for invaluable help with technical and administrative aspects of this work. Thanks to Maria Littmann for developing the project idea and reviewing the first text draft. Thanks to the entire Rostlab (TUM) for their support and productive discussions. Thanks to Svenja Lemke for helpful comments on the manuscript. Thanks to the MTA-ELTE Momentum Bioinformatics Research Group and the Kurgan Lab for making their respective prediction methods IUPred2A and DeepDISOBind publicly available. Last but not least thanks to the providers of MobiDB (Silvio CE Tosatto and Damiano Piovesan), the experimentalist who make their precious data available, and all developers who make their methods freely available.

## Author contributions

L.R.J. implemented and evaluated the method and co-wrote the manuscript. C.M. supervised the project, performed analyses, and co-wrote the manuscript. M.H. co-conceived and supervised the project. B.R. supervised and guided the work. All authors read and approved the final manuscript.

## Additional Information

This work was supported by the Bavarian Ministry of Education through funding to the TUM, by a grant from the Alexander von Humboldt foundation through the German Ministry for Research and Education (BMBF: Bundesministerium für Bildung und Forschung), and by a grant from Deutsche Forschungsgemeinschaft (DFG-GZ: RO1320/4-1). The authors declare no conflicts of interest.

