## Supplementary Material for "Protein Embeddings Predict Binding Residues in Disordered Regions"

### Table of Contents for Supporting Online Material

### Short description of Supporting Online Material

Figures SOM_F1 – SOM_F4 display ROC (receiver operating characteristic) curves of all trained models, the models' performance on the validation set, the training curve of each model's best cross validation (CV)-fold and their precision-recall curve, respectively. The second plot SOM_F2 was used for deciding on the final predictor. Figures SOM_F5 – SOM_F7 explore IDBindT5's predictions on the test set in terms of binding region continuity and a prediction profile. Figure SOM_F8 depicts the feature weights of the model's input layer. Table SOM_T1 lists the number of proteins and residues of each data set. Table

SOM_T2 provides the parameters for training the models. Tables SOM_T3 and SOM_T4 contain all performance assessments that were created for the models. Table SOM_T5 shows the test set performance of two disorder predictors, which were used as input to IDBindT5 instead of experimental annotation (see Table SOM_T4). Table SOM_T6 contains performance assessments on the CAID2 binding benchmark set. Table SOM_T7 gives the results of Welch's t-test applied to our final model and two state-of-the-art methods. Lastly, Table SOM_T8 provides runtimes of IDBindT5 and SOTA models under different conditions.

### Material

**Supplementary Fig. SOM_F1: ROC curves on the validation set.**

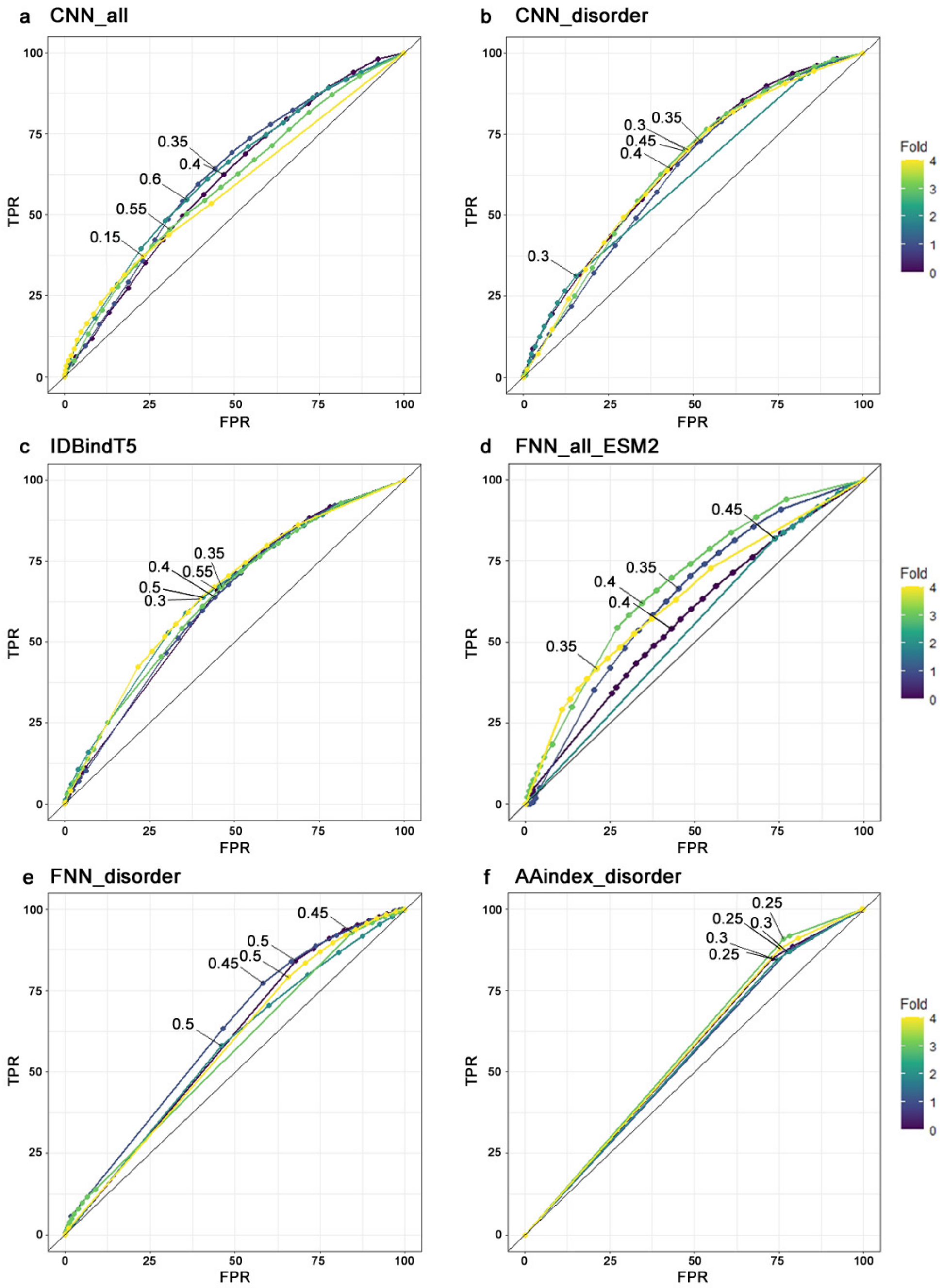

**Supplementary Fig. SOM_F1: ROC curves on the validation set.** These ROC curves were used to determine the optimal cut off for classifying a residue as binding (output probability of the model > cut off) or non-binding (output probability ≤ cut off). The aim was to achieve a good precision while avoiding a large drop in recall. The chosen cut-offs are annotated for each model and CV-fold.

**Supplementary Fig. SOM_F2: Performance on the validation set.**

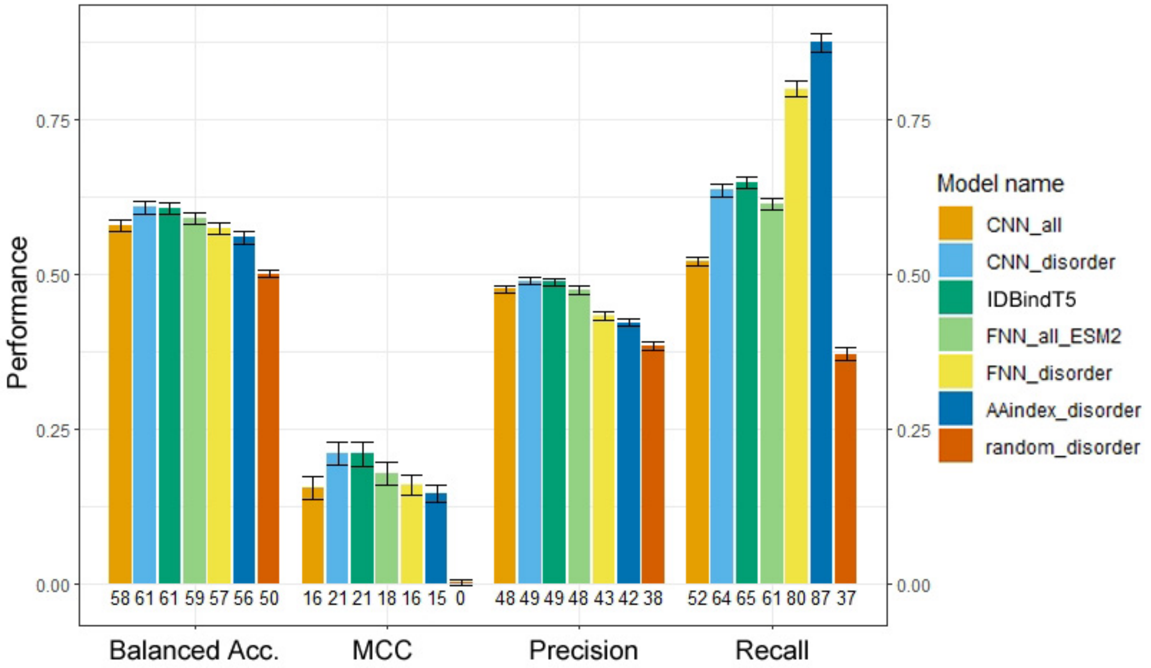

**Supplementary Fig. SOM_F2: Performance on the validation set.** Average performance and standard error of the six most relevant predictors and a random baseline predictor for binding regions on the validation set. The random predictor only considers the class imbalance in the disordered training set for its predictions. The performance is captured by the four per-residue measures: balanced accuracy (Eqn. 5), MCC (Eqn. 7), precision (Eqn. 1) and recall (Eqn. 2) (definitions see *Methods*). The numbers below each performance bar show the respective performance times 100, rounded to the closest integer. All models performed significantly better than the random baseline in all aspects, but they suffer from low precision. *FNN_all_ESM2*, which applies the *IDBINDT5* architecture to ESM-2 [1] embeddings, is significantly worse than model *IDBINDT5* in the aspects of precision and recall. This suggests that ProtT5 might be better suited for predicting disordered residue features than ESM-2. The simplistic model *AAindex_disorder*, which represents a baseline for models based on AA propensities only, can compete with *CNN_all* and *FNN_disorder*, but is outperformed by *CNN_disorder*, *IDBINDT5* and *FNN_all_ESM2*. We cannot see a trend in performance that would lead us to favor one architecture or training residue subset over another. However, based on predictive performance and the observation that our FNNs are faster than our CNNs, *IDBINDT5* was chosen as final predictor. More performance measures and exact values can be found in Table SOM_T3.

**Supplementary Fig. SOM_F3: Training curves of all models.**

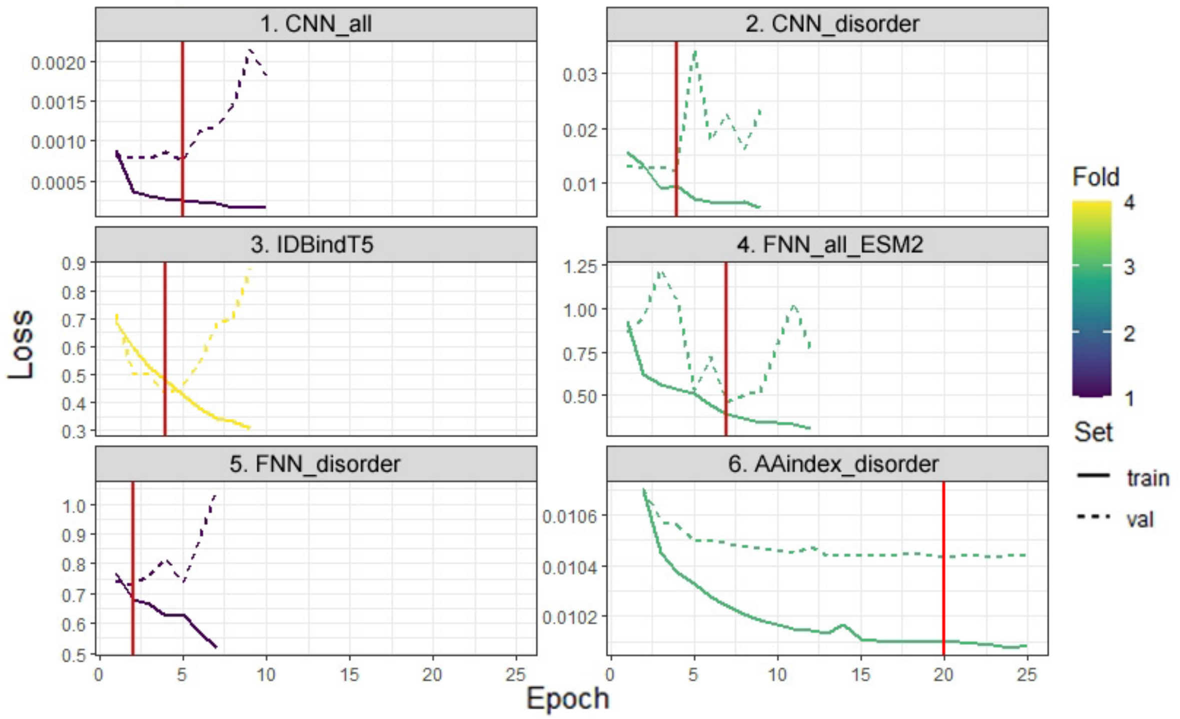

**Supplementary Fig. SOM_F3: Training curves of all models.** Loss on training and validation set, monitored for each epoch during training of the best CV-folds of the six relevant models. The red vertical line indicates which version was saved via early stopping with patience of five epochs. The final model, *IDBindT5*, trained for only nine epochs before early stopping saved the model of epoch four. All models trained on protein embeddings only train for a short while, whereas the model based on biochemical and -physical features, *AAindex_disorder*, trains for a much longer time of 25 epochs. We suspect that this difference might be caused by the rich information content of the embeddings, meaning the ML model only needed to filter for a few relevant features and weigh them accordingly. This has also been observed for other models trained on ProtT5 embeddings, such as bindEmbed21DL [2], TMbed [3], VESPA [4] and EMBER2 [5].

**Supplementary Fig. SOM_F4: Precision and recall at different cutoffs.**

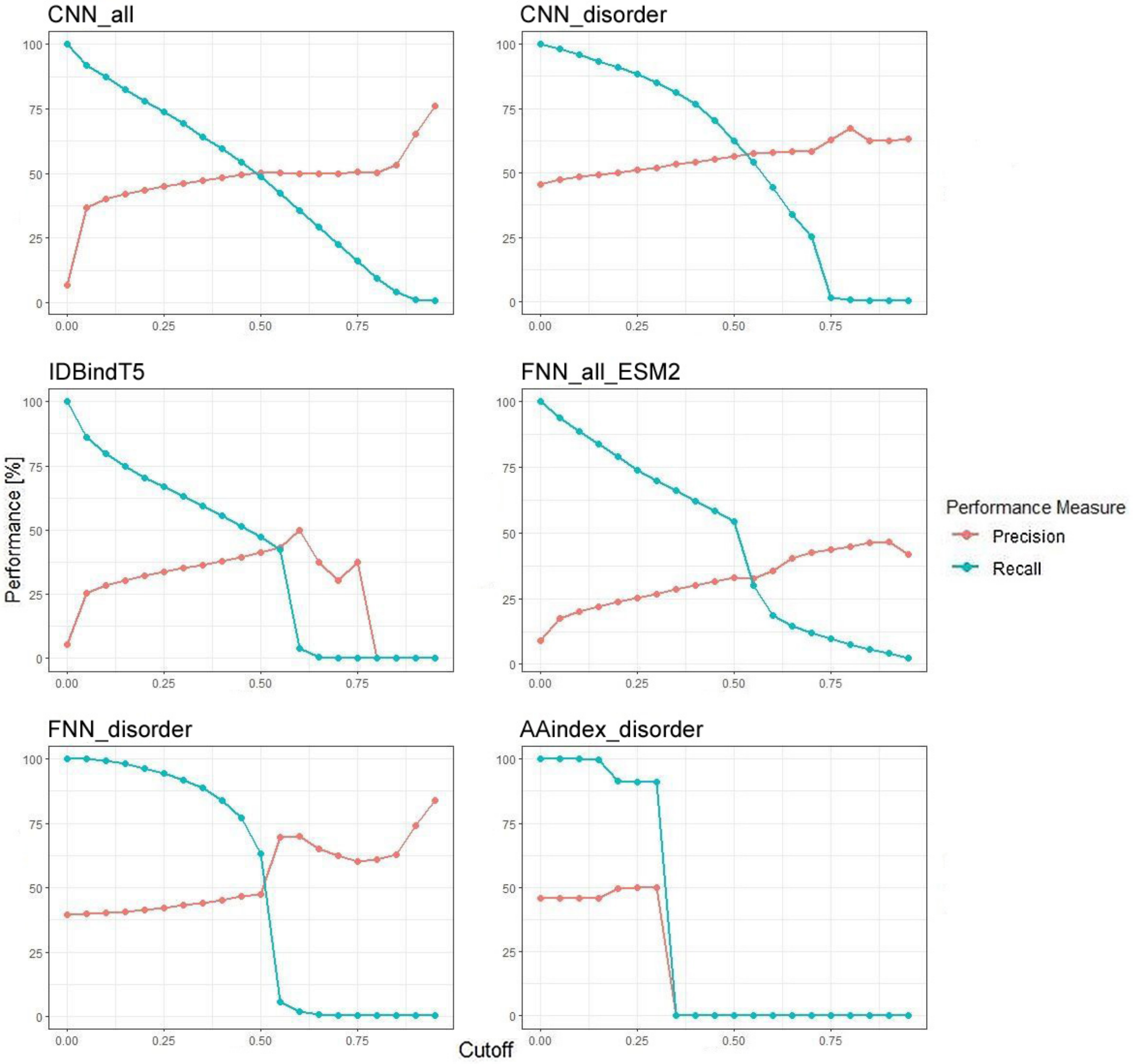

**Supplementary Fig. SOM_F4: Precision and recall at different cut offs.** Precision (Eqn. 1) and recall (Eqn. 2) change across different cut offs for the classification as binding or non-binding on the validation set. The x-axis gives the raw output of the model for a prediction, a.k.a. the prediction strength. The y-axis gives the average performance at the respective cut off. All curves are cumulative, i.e., the metric of all residues predicted with a value ≥ cut off is as shown on the y-axis. High cut offs correspond to more reliable binding predictions and lower cut offs lead to more precise non-binding predictions. For the optimal balanced performance, we chose *IDBindT5* at cut off 0.3 as final model. However, if a large dataset shall be scanned for few, but higher-confidence binding predictions, we recommend using *FNN_disorder* at cut off 0.55, instead.

**Supplementary Fig. SOM_F5: Length distribution of binding regions.**

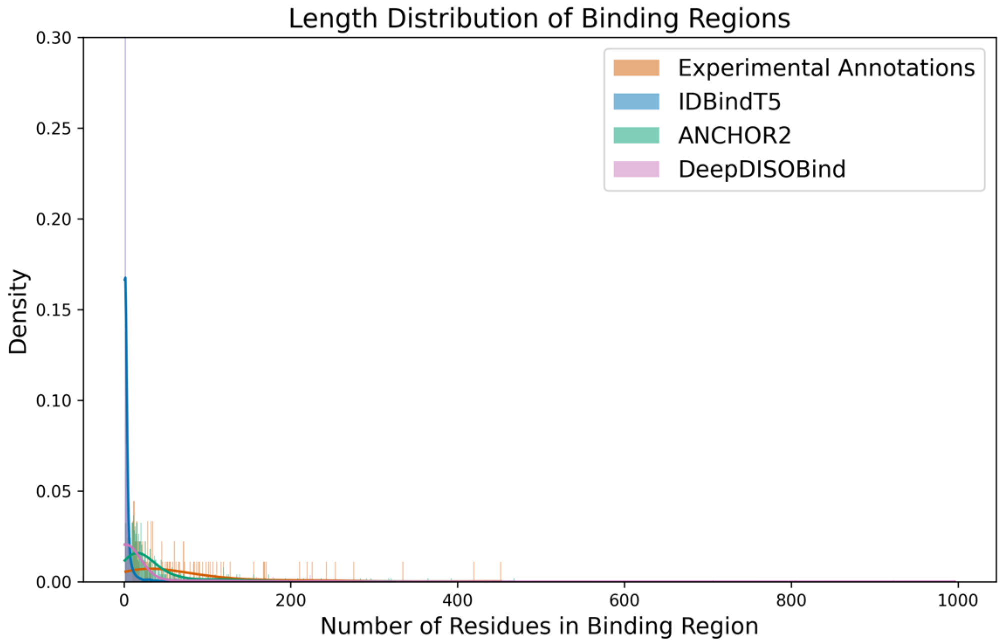

**Supplementary Fig. SOM_F5: Length distribution of binding regions.** The histogram shows the distribution of lengths of binding regions in for experimental and predicted values of the test set with Experimental Annotations in orange, IDBindT5 in blue, ANCHOR2 [6] in green, and DeepDISOBind [7] in pink. The x-axis represents the number of residues in the binding region, while the y-axis represents the density of occurrences. Each dataset represents the lengths of stretches of binding residues, where a value of 1 indicates a binding residue and 0 indicates a non-binding residue. Binding regions are identified as contiguous stretches of residues labelled as binding (1).

**Supplementary Fig. SOM_F6: Length distribution of binding regions.**

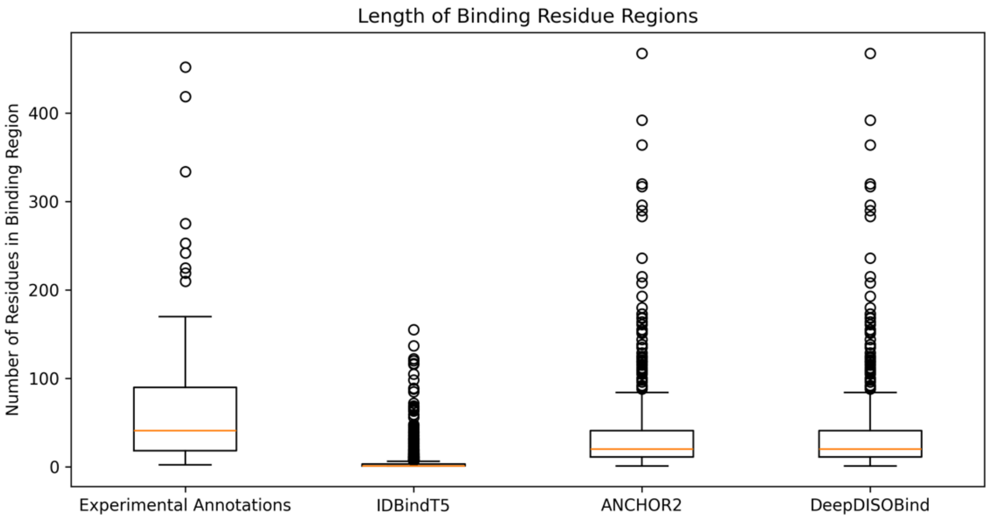

**Supplementary Fig. SOM_F6: Length distribution of binding regions.** The box plot depicts the distribution of binding residue region lengths in the test set for experimental annotations, IDBindT5, ANCHOR2 [6], and DeepDISOBind [7] predictions (from left to right). Each boxplot comprises a box representing the interquartile range (IQR) with the median marked by a horizontal line within the box, whiskers extending to the minimum and maximum values within 1.5 times the IQR from the quartiles, and individual data points beyond the whiskers indicating outliers. The x-axis labels specify the source of the annotations, while the y-axis quantifies the number of residues in the binding regions. This visualization facilitates comparisons between the datasets regarding measures of central tendency, spread, and potential anomalies.

**Supplementary Fig. SOM_F7: Prediction Profile of IDBindT5 for test set.**

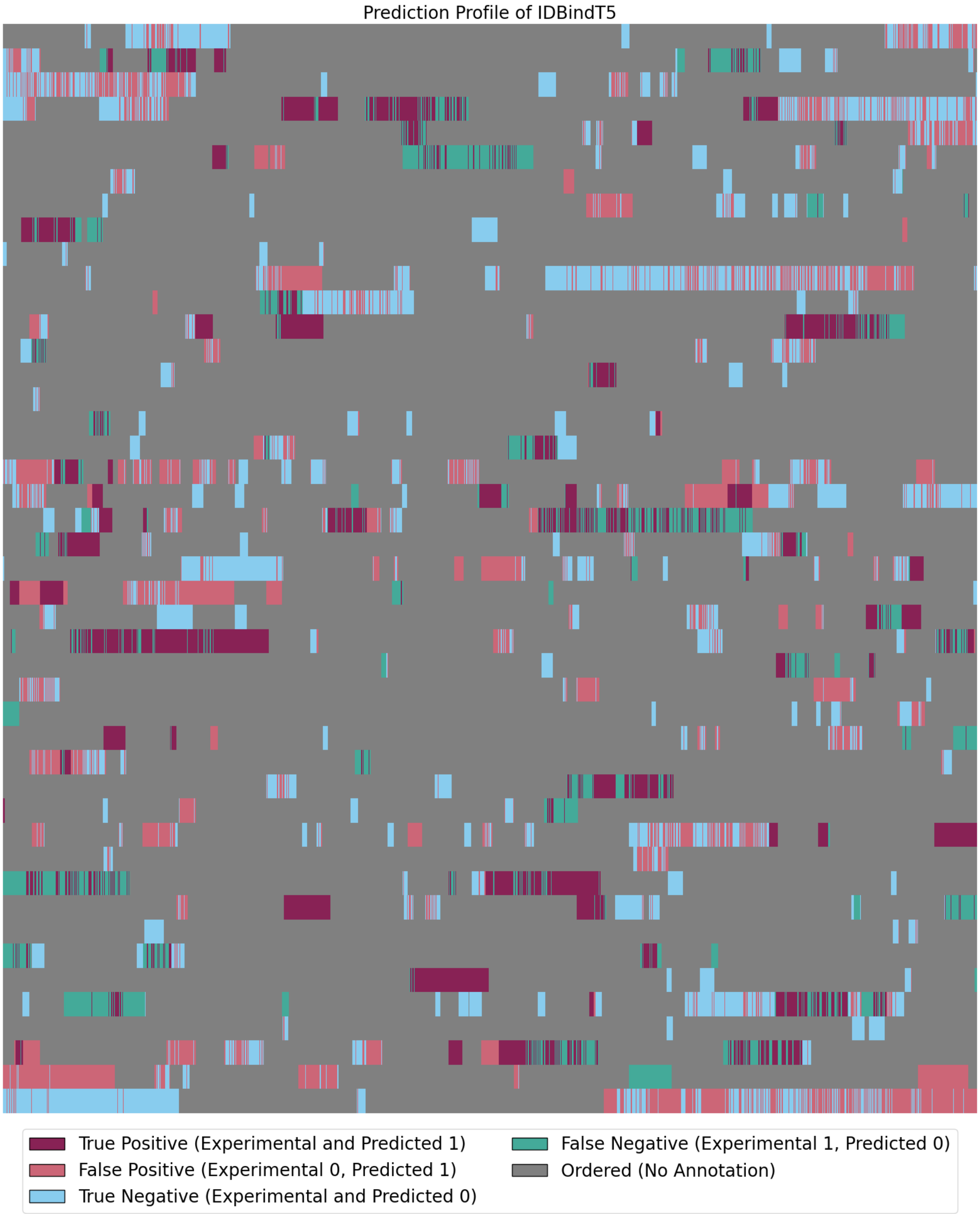

**Supplementary Fig. SOM_F7: Prediction Profile of IDBindT5 for test set.** The prediction profile for IDBindT5 shows all 92,373 of 195 proteins in the test set concatenated. Each pixel is colored by one of the following labels: "True Positive" with both experimental and predicted labeled binding in dark red, "False Positive" with experimental labeled non-binding and predicted labeled binding in light red, “True Negative” with both experimental and predicted labeled non-binding in blue, “False Negative” with experimental labeled binding and predicted labeled non-binding in green, or lastly ”Ordered” residues with no annotation in grey.

**Supplementary Fig. SOM_F8: Feature Importance of IDBindT5.**

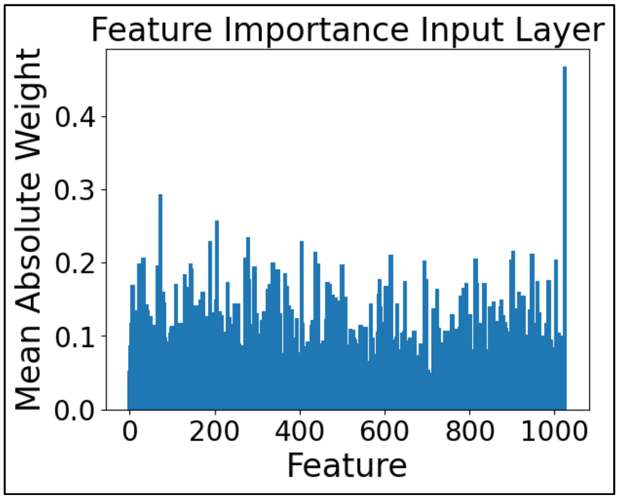

**Supplementary Fig. SOM_F8: Feature Importance of IDBindT5.** IDBindT5 is a feedforward neural network with one hidden layer taking a vector of length 1025 as input (ProtT5 embeddings [5] length 1024 and binary (dis)order annotation). The plot shows the feature importance as mean absolute weight of the input layer. The last layer with disorder annotation has the highest mean absolute weight, while the pLM embedding layer weights are more evenly distributed.

**Supplementary Table SOM_T1: Composition of the final data set.** *

| Set | Subset of residues | Binding proteins | Non-binding proteins | Binding residues | Non-binding residues |
|---|---|---|---|---|---|
| **Training Set** | All | 697 | 1,083 | 64,228 | 896,569 |
| | IDPRs only | | | | 101,381 |
| CV Set 0 | All | 139 | 223 | 10,235 | 165,230 |
| | IDPRs only | | | | 20,008 |
| CV Set 1 | All | 153 | 214 | 14,248 | 193,846 |
| | IDPRs only | | | | 21,698 |
| CV Set 2 | All | 140 | 196 | 10,386 | 155,712 |
| | IDPRs only | | | | 18,665 |
| CV Set 3 | All | 143 | 225 | 17,906 | 186,048 |
| | IDPRs only | | | | 21,390 |
| CV Set 4 | All | 122 | 225 | 11,453 | 195,733 |
| | IDPRs only | | | | 19,620 |
| **Test Set** | All | 80 | 115 | 6,657 | 85,716 |
| | IDPRs only | | | | 11,644 |

* Number of proteins and residues in their respective data sets after redundancy reduction, as described in *Methods - Data set*. The rows *CV Set 0 – 4* represent the five distinct cross validation subsets of the training set. The FASTA files can be found in the Github repository (https://github.com/jahnl/binding_in_disorder/tree/main/dataset/MobiDB_dataset_2).

**Supplementary Table SOM_T2: Model parameters.** *

| Parameter | 1. CNN_all | 2. CNN_disorder | 3. IDBindT5 |
|---|---|---|---|
| **Input** | ProtT5 embeddings + disorder feature | ProtT5 embeddings | ProtT5 embeddings + disorder feature |
| **Balancing method** | Oversampling positive proteins | Undersampling negative proteins | Undersampling negative residues ** |
| **Residue subset** | All | IDPRs only | All |
| **Architecture** | CNN | CNN | FNN |
| **# Layers** | 5 | 5 | 3 |
| **Kernel size** | 5 | 5 | - |
| **Dropout** | 0.0 | 0.0 | 0.0 |
| **Batch size** | 1 protein | 1 protein | 512 residues |
| **Learning rate** | 0.002 | 0.002 | 0.01 |
| **Patience** | 5 | 5 | 5 |

| Parameter | 4. FNN_all_ESM2 | 5. FNN_disorder | 6. AAindex_disorder |
|---|---|---|---|
| **Input** | ESM-2 embeddings + disorder feature | ProtT5 Embeddings | AAindex1 |
| **Balancing method** | Undersampling negative residues ** | Undersampling negative residues ** | Undersampling negative proteins |
| **Residue subset** | All | IDPRs only | IDPRs only |
| **Architecture** | FNN | FNN | CNN |
| **# Layers** | 3 | 3 | 5 |
| **Kernel size** | - | - | 5 |
| **Dropout** | 0.0 | 0.0 | 0.0 |
| **Batch size** | 512 residues | 512 residues | 1 protein |
| **Learning rate** | 0.01 | 0.01 | 0.002 |
| **Patience** | 5 | 5 | 5 |

* Parameters of the most relevant models. Models 1-3 and 5 perform best on the validation set in their respective category, which are: CNNs on all residues, FNNs on all residues, CNNs on disordered residues only, FNNs on disordered residues only. Model 5 is trained on ESM-2 [1] instead of ProtT5 [8] embeddings and applies the parameters of the *IDBindT5* model. The last model is trained on AAindex1 [9] features and adapts the parameters of the *CNN_disorder* model. An AAindex-model trained on all residues

was tried out as well, however, it did not learn anything useful for binding region prediction.

** Undersampling negative residues describes a balancing method, where negative residues within and outside of IDPRs are undersampled with different ratios, so that each subset reaches the same abundance as the positive residues.

---

**Supplementary Table SOM_T3: Performance assessment on the validation set.** *

| Model | 1. CNN_all | 2. CNN_disorder | 3. IDBindT5 |
|---|---|---|---|
| **Precision** | 47.7 ± 0.6% | **48.9 ± 0.6%** | **48.9 ± 0.6%** |
| **Recall** | 52.1 ± 0.8% | 63.6 ± 0.9% | 64.8 ± 1.0% |
| **Neg. Precision** | 67.7 ± 0.9% | 71.6 ± 0.9% | 71.6 ± 0.8% |
| **Neg. Recall** | **63.6 ± 0.8%** | 58.0 ± 0.9% | 56.7 ± 0.8% |
| **Balanced Acc.** | 57.9 ± 0.9% | **60.8 ± 0.9%** | **60.8 ± 0.9%** |
| **F1 Score** | 0.498 ± 0.009 | 0.553 ± 0.009 | 0.557 ± 0.009 |
| **MCC** | 0.156 ± 0.019 | **0.211 ± 0.018** | 0.210 ± 0.019 |

| Model | 4. FNN_all_ESM2 | 5. FNN_disorder | 6. AAindex_disorder | Random_disorder |
|---|---|---|---|---|
| **Precision** | 47.5 ± 0.6% | 43.3 ± 0.7% | 42.3 ± 0.6% | 38.5 ± 0.7% |
| **Recall** | 61.4 ± 0.8% | 80.0 ± 1.3% | **87.5 ± 1.5%** | 37.2 ± 1.1% |
| **Neg. Precision** | 69.9 ± 0.9% | 73.7 ± 0.9% | **75.6 ± 1.2%** | 61.7 ± 0.7% |
| **Neg. Recall** | 56.9 ± 0.8% | 35.0 ± 0.8% | 24.5 ± 1.0% | 63.1 ± 0.9% |
| **Balanced Acc.** | 59.2 ± 1.0% | 57.5 ± 0.9% | 56.0 ± 1.0% | 50.1 ± 0.6% |
| **F1 Score** | 0.536 ± 0.008 | 0.562 ± 0.009 | **0.570 ± 0.010** | 0.378 ± 0.006 |
| **MCC** | 0.179 ± 0.019 | 0.160 ± 0.016 | 0.146 ± 0.014 | 0.002 ± 0.004 |

* Performance assessment of the most relevant predictors in comparison to each other and a random baseline on the IDPRs in the validation set. The performance measures were calculated according to the definition in the *Methods* chapter (Eqn. 1-8) and rounded to three decimal points. The number after the ± refers to the 95% confidence interval of the standard error. The numerically best performances per metric are printed in bold.

**Supplementary Table SOM_T4: Performance assessment on the test set.** *

| Model | 1. CNN_all | 2. CNN_ disorder | 3. IDBindT5 | 4. FNN_all_ ESM2 | 5. FNN_ disorder |
|---|---|---|---|---|---|
| **Precision** | 42.2 ± 1.9% | 40.1 ± 2.1% | **43.2 ± 1.9%** | 36.9 ± 2.5% | 39.5 ± 2.1% |
| **Recall** | 61.0 ± 3.3% | 64.8 ± 3.0% | 58.7 ± 2.6% | 48.4 ± 2.5% | 72.5 ± 3.5% |
| **Neg. Precision** | 70.1 ± 2.7% | 69.0 ± 2.9% | 70.2 ± 2.8% | 64.1 ± 2.6% | 68.6 ± 2.9% |
| **Neg. Recall** | 52.4 ± 2.4% | 44.7 ± 2.5% | 55.8 ± 2.5% | 52.7 ± 2.2% | 35.1 ± 2.5% |
| **Balanced Acc.** | 56.7 ± 2.8% | 54.8 ± 2.7% | 57.2 ± 3.6% | 50.6 ± 2.8% | 53.8 ± 2.6% |
| **F1 Score** | 0.499 ± 0.026 | 0.496 ± 0.027 | 0.498 ± 0.027 | 0.419 ± 0.028 | 0.511 ± 0.030 |
| **MCC** | 0.128 ± 0.053 | 0.093 ± 0.053 | 0.139 ± 0.070 | 0.011 ± 0.055 | 0.078 ± 0.054 |

| Model | Consensus of models 2. and 3. | 6. AAindex_ disorder | Random_ disorder | ANCHOR2 [6] | DeepDISO-Bind [7] |
|---|---|---|---|---|---|
| **Precision** | 43.1 ± 1.8% | 39.0 ± 2.0% | 36.3 ± 2.3% | 39.5 ± 1.6% | 42.2 ± 2.4% |
| **Recall** | 60.6 ± 2.8% | **86.1 ± 5.0%** | 36.2 ± 2.2% | 45.0 ± 2.6% | 51.4 ± 3.2% |
| **Neg. Precision** | 70.6 ± 2.9% | **74.3 ± 3.6%** | 62.8 ± 2.4% | 65.2 ± 2.7% | 70.6 ± 3.3% |
| **Neg. Recall** | 54.2 ± 2.3% | 22.9 ± 3.0% | 62.9 ± 3.4% | 59.9 ± 2.4% | 62.3 ± 3.4% |
| **Balanced Acc.** | **57.4 ± 3.0%** | 54.5 ± 2.8% | 49.6 ± 2.2% | 52.4 ± 2.7% | 56.9 ± 5.6% |
| **F1 Score** | 0.503 ± 0.030 | **0.537 ± 0.033** | 0.363 ± 0.021 | 0.421 ± 0.036 | 0.463 ± 0.045 |
| **MCC** | **0.142 ± 0.065** | 0.110 ± 0.036 | -0.009 ± 0.019 | 0.048 ± 0.060 | 0.133 ± 0.120 |

* Performance assessment on the IDPRs in the test set. The most relevant predictors are compared to each other, a random baseline, the established tools ANCHOR2 [6] and DeepDISOBind [7], and IDBindT5 on several predicted disorder labels. The performance measures were calculated according to the definition in the *Methods* chapter (Eqn. 1-8) and rounded to three decimal points. The number after the ± refers to the 95% confidence interval of the standard error. The numerically best performances per metric in bold.

#### Supplementary Table SOM_T5: IDBindT5 performance inputting predicted disorder*

| Disorder Predictor | SETH [10] | Alphafold-disorder [11] |
|---|---|---|
| **Precision** | 37.3% | 40.2% |
| **Recall** | 66.5% | 67.3% |
| **Neg. Precision** | 89.7% | 94.9% |
| **Neg. Recall** | 72.3% | 85.8% |
| **Balanced Acc.** | 69.4% | 76.6% |
| **F1 Score** | 0.477 | 0.503 |
| **MCC** | 0.324 | 0.432 |

| Binding site predictor | IDBindT5 with SETH disorder predictions | IDBindT5 with AlphaFold2 disorder predictions | IDBindT5 with experimental disorder annotations |
|---|---|---|---|
| **Precision** | 40.9 ± 2.0% | 42.1 ± 1.9% | **43.2 ± 1.9%** |
| **Recall** | 39.5 ± 3.0% | 42.3 ± 2.8% | **58.7 ± 2.6%** |
| **Neg. Precision** | 66.1 ± 2.4% | 66.9 ± 2.6% | **70.2 ± 2.8%** |
| **Neg. Recall** | **67.3 ± 2.8%** | 66.8 ± 2.5% | 55.8 ± 2.5% |
| **Balanced Acc.** | 53.4 ± 3.2% | 54.5 ± 3.2% | **57.2 ± 3.6%** |
| **F1 Score** | 0.402 ± 0.026 | 0.422 ± 0.028 | **0.498 ± 0.027** |
| **MCC** | 0.069 ± 0.066 | 0.091 ± 0.065 | **0.139 ± 0.070** |

* We applied pre-existing disorder predictors to our test set to investigate the impact of using predicted instead of curated disorder information in the input data to IDBindT5. The upper table shows the numeric performance of these disorder predictors on the test set, the lower one shows IDBindT5's performance given these disorder labels. The performance measures were calculated according to the definition in the *Methods* chapter (Eqn. 1-8) and rounded to three decimal points.

**Supplementary Table SOM_T6: Performance of disorder predictors on CAID2's binding dataset** *

| Model | IDBindT5 | ANCHOR2 [6] | DeepDISOBind [7] | Random_disorder |
|---|---|---|---|---|
| **Residue Subset** | **IDPRs only** | **IDPRs only** | **IDPRs only** | **IDPRs only** |
| **Precision** | 61.4 ± 2.7% | 57.0 ± 2.5% | **61.5 ± 3.2%** | 55.9 ± 2.5% |
| **Recall** | 69.0 ± 3.6% | 60.6 ± 3.2% | **69.4 ± 4.0%** | 36.5 ± 1.1% |
| **Neg. Precision** | 51.0 ± 1.9% | 43.1 ± 2.4% | **51.3 ± 3.0%** | 42.4 ± 2.5% |
| **Neg. Recall** | 42.6 ± 3.8% | 39.4 ± 3.1% | 42.6 ± 2.4% | **61.9 ± 4.4%** |
| **Balanced Acc.** | 55.8 ± 4.5% | 50.0 ± 4.5% | **56.0 ± 4.0%** | 49.2 ± 2.7% |
| **F1 Score** | 0.650 ± 0.028 | 0.587 ± 0.025 | **0.652 ± 0.046** | 0.441 ± 0.020 |
| **MCC** | 0.120 ± 0.086 | 0.001 ± 0.093 | **0.124 ± 0.084** | -0.017 ± 0.024 |

| Model | IDBindT5 | ANCHOR2 | DeepDISOBind | Random_disorder |
|---|---|---|---|---|
| **Residue Subset** | **All** | **All** | **All** | **All** |
| **Precision** | 37.6 ± 2.3% | 17.0 ± 3.5% | 17.5 ± 3.1% | **57.2 ± 2.5%** |
| **Recall** | 69.0 ± 3.4% | 60.6 ± 2.4% | **69.4 ± 2.4%** | 37.0 ± 1.6% |
| **Neg. Precision** | **96.5 ± 3.1%** | 94.6 ± 3.1% | 95.5 ± 3.4% | 43.2 ± 2.4% |
| **Neg. Recall** | **88.3 ± 2.2%** | 69.9 ± 1.4% | 66.6 ± 1.3% | 63.4 ± 1.2% |
| **Balanced Acc.** | **78.6 ± 1.6%** | 65.2 ± 3.0% | 68.0 ± 3.3% | 50.2 ± 1.1% |
| **F1 Score** | **0.487 ± 0.023** | 0.266 ± 0.027 | 0.280 ± 0.034 | 0.449 ± 0.020 |
| **MCC** | **0.442 ± 0.034** | 0.188 ± 0.061 | 0.217 ± 0.078 | 0.004 ± 0.020 |

* Performance assessment on CAID2's [12] binding dataset. The performance was computed for both the IDPR subset (naturally including all positive datapoints), like we did it for our test set in this project, and all residues, like it was done in the CAID challenge [12]. The performance measures were calculated according to the definition in the *Methods* chapter (Eqn. 1-8) and rounded to three decimal points. The number after the ± refers to the 95% confidence interval of the standard error. The numerically best performances per metric and residue subset are printed in bold.

**Supplementary Table SOM_T7: Welch's t-test applied to models on the test set.** *

| Balanced Accuracy | | | | |
|---|---|---|---|---|
| **t-statistic \ p-value** | **IDBindT5** | **Random_disorder** | **ANCHOR2 [6]** | **DeepDISOBind [7]** |
| **IDBindT5** | \ | **0.006** | 0.571 | 0.610 |
| **Random_disorder** | **- 2.790** | \ | **0.036** | 0.418 |
| **ANCHOR2** | - 0.567 | **2.124** | \ | 0.859 |
| **DeepDISOBind** | - 0.514 | 0.822 | - 0.178 | \ |

| MCC | | | | |
|---|---|---|---|---|
| **t-statistic \ p-value** | **IDBindT5** | **Random_disorder** | **ANCHOR2** | **DeepDISOBind** |
| **IDBindT5** | \ | **0.001** | 0.854 | 0.642 |
| **Random_disorder** | **- 3.460** | \ | **0.008** | 0.290 |
| **ANCHOR2** | - 0.184 | **2.729** | \ | 0.735 |
| **DeepDISOBind** | - 0.469 | 1.082 | - 0.341 | \ |

| Precision | | | | |
|---|---|---|---|---|
| **t-statistic \ p-value** | **IDBindT5** | **Random_disorder** | **ANCHOR2** | **DeepDISOBind** |
| **IDBindT5** | \ | 0.307 | 0.897 | 0.814 |
| **Random_disorder** | - 1.024 | \ | 0.286 | 0.582 |
| **ANCHOR2** | 0.130 | 1.069 | \ | 0.743 |
| **DeepDISOBind** | - 0.235 | 0.552 | - 0.645 | \ |

| Recall | | | | |
|---|---|---|---|---|
| **t-statistic \ p-value** | **IDBindT5** | **Random_disorder** | **ANCHOR2** | **DeepDISOBind** |
| **IDBindT5** | \ | **<0.001** | **0.013** | 0.244 |
| **Random_disorder** | **- 5.038** | \ | 0.210 | 0.074 |
| **ANCHOR2** | **- 2.508** | 1.259 | \ | 0.440 |
| **DeepDISOBind** | - 1.171 | 1.817 | 0.774 | \ |

* The scipy implementation of Welch's t-test [13, 14] was applied to different metrics of all models on the test set. In contrast to the student's t-test it does not assume equal population variance. If the p-value (upper-right triangles) is smaller than 0.05, then we have evidence against the null hypothesis of equal population means [13], so there is a

significant difference (printed in bold). The t-statistic (lower-left triangles) quantifies the difference between the arithmetic means of the rows's model vs. the columns's model. All values are rounded to three decimal points.

**Supplementary Table SOM_T8: Speed assessment ***

| **Conditions** | | | **Predictor (median runtime)** |
|---|---|---|---|
| **Dataset** | **Machine** | **Use of GPU** | **IDBindT5** |
| P09938 (399 AAs) | Consumer-grade | No | 2.246 s |
| P09938 (399 AAs) | Consumer-grade | Yes | 2.378 s |
| Test set (Mobi195) | Consumer-grade | No | 12.594 s |
| Test set (Mobi195) | Consumer-grade | Yes | 8.146 s |

| **Conditions** | | | **Predictor (absolute runtime)** | | |
|---|---|---|---|---|---|
| **Dataset** | **Machine** | **Use of GPU** | **IDBindT5** | **ANCHOR2 [6]** | **eepDISOBind [7]** |
| Mobi11k | Cluster | Yes | 2 h, 10 min | 21 min | 33 h, 22 min |

* Speed assessment under different conditions (dataset, machine, usage of GPU). We provide the median runtime of 10 single prediction runs on the small datasets (upper table) per condition, and the absolute runtimes on the biggest dataset (lower table). The sequence of the protein P09938 (Ribonucleoside-diphosphate reductase small chain 1) has a sequence length of 399, which matches the median sequence length of our training set (Mobi2k). The consumer-grade machine's stats are: CPU: Intel Core i7, 16 GB RAM, 4 cores; GPU: NVIDIA GeForce MX350, 9.9 GB vRAM. The cluster's stats are: CPU: 2x Intel Xeon Gold 6248, 400 GB RAM, 20 cores; GPU: 4x Quadro RTX 8000, 46 GB vRAM.
